# Oncoprotein GT198 Vaccination Delays Tumor Growth in MMTV-PyMT Mice

**DOI:** 10.1101/693606

**Authors:** Bhagelu R. Achyut, Hao Zhang, Kartik Angara, Nahid F. Mivechi, Ali S. Arbab, Lan Ko

## Abstract

Different effects of anticancer drugs between mouse and human have caused increasing concerns. A better understanding of cancer initiation between the two species is needed. We have previously identified an oncoprotein GT198 (*PSMC3IP*) in human breast cancer. In this report, we investigated GT198 in MMTV-PyMT mouse mammary gland tumors and found a reconcilable mechanism in human and mouse. Specifically, distinct tumor initiating stimuli in human and mouse result in a common GT198-mediated tumorigenic pathway in both species. Here we show, similar to human breast cancer even before a tumor appears, GT198 has overexpressed in mouse tumor stroma including pericyte stem cells, descendent adipocytes, fibroblasts, and myoepithelial cells. Using recombinant GT198 protein as an antigen, we vaccinated MMTV-PyMT mice and found that the GT198 vaccine delayed mouse tumor growth and reduced lung metastasis. The antitumor effects in vaccinated mice were linearly correlated with serum titers of GT198 antibody, which can recognize cell surface GT198 protein on viable tumor cells confirmed by FACS. Furthermore, tumor cells isolated from MMTV-PyMT mice were re-implanted into normal FVB/N mice, GT198^+^ tumor cells induced faster tumor growths than GT198^-^ tumor cells. Together, this first study of GT198 vaccine in mouse showed its effectiveness in antitumor and anti-metastasis. The finding may accelerate future development of GT198 immunotherapy in human cancer. Our finding also indicates that even though distinct cancer-initiation stimuli exist between mouse and human, a common tumorigenic pathway mediated by oncoprotein GT198 is shared in both species.

## INTRODUCTION

In 1953, the mouse polyomavirus was isolated as one of the first DNA viruses with its potential role in tumorigenesis (1,2). Polyomavirus was found to induce tumors when inoculated into a variety of mouse tissues. Subsequently, it was established that polyomavirus expresses several tumor antigens, including the middle T antigen, which is necessary and sufficient for polyomavirus-induced tumor transformation (3-5).

The polyomavirus middle T antigen is a membrane-bound protein containing 421 amino acids with multiple phosphorylation sites. It does not have kinase activity itself but serves as a scaffold to interact and activate a panel of kinases and phosphatases, including but not limited to, Src family tyrosine kinases, protein phosphatase 2A, PI3 kinase, Shc and Ras, which in turn stimulate cellular signal transduction pathways (2,4). In particular, the Ras pathway with downstream mitogen-activated protein (MAP) kinase activation has been extensively studied for tumorigenesis in molecular or cellular contexts, and in mouse tumor models (3,4,6). Because complex nuclear protein factors are targeted by virus-induced kinase signaling, the nuclear mechanism responsible for tumor initiation remains undetermined.

The effects of middle T antigen in mouse mammary glands were investigated in transgenic mice carrying mouse mammary tumor virus long terminal repeat (MMTV-PyMT), where the middle T antigen can be specifically expressed in mammary glands under the MMTV promoter. This resulted in widespread, multifocal, and spontaneous mammary tumor formation with onset as early as 3-4 weeks of age (7). Similarly, mammary gland-specific expression of v-Ha-ras oncogene in MMTV-Ras transgenic mice also showed widespread and spontaneous mammary gland tumors (8). Tumor metastasis is often found in mouse lung and lymph nodes.

Despite substantial evidence of tumor formation in mouse models, the obvious difference may exist between mouse and human tumor initiation, which become an increasing concern (9-11). Since the goal of cancer research is the development of therapies for human with drug target identification, it was a challenge to reconcile, for example, human breast cancer versus mouse mammary gland tumor, in their physiology, pathology, and molecular signaling.

We have previously identified a human breast cancer initiating oncogene called *GT198* (gene symbol *PSMC3IP*) (12). In this study, we investigate oncoprotein GT198 expression in MMTV-PyMT and MMTV-Ras mice and carry out vaccination using GT198 protein as an antigen. We found that the differences between human and mouse can be reconciled. Specifically, distinct tumor initiating stimuli in human and mouse resulted in a common GT198-mediated pathway of tumor development in both species.

GT198 is a DNA-binding protein dimer, containing 217 amino acids in its monomer (13). GT198 binds to single- and double-stranded DNAs and the binding is not sequence-specific. GT198 is a transcriptional coactivator stimulating nuclear receptor-mediated gene activation (13,14). GT198 protein is MAP kinase phosphorylation regulated (12,13). GT198 is also known as Hop2/TBPIP, acts as a crucial DNA repair factor by participating in homologous DNA recombination and regulating meiosis (15-17).

The human *GT198* gene is located at chromosome 17q21, 470 Kb proximal to BRCA1, a hot cancer locus previously linked to breast and ovarian cancer predisposition. Germline mutations in *GT198* have been identified in familial and early-onset human breast and ovarian cancer (18,19), as well as in familial ovarian disease (20). Somatic mutations in *GT198* are prevalent and recurrent in sporadic breast, ovarian, prostate, uterine, and fallopian tube cancers, where mutant GT198 leads to constitutive activation in transcription (18,21).

GT198 regulates mouse stem cells at the embryoid body stage (21). In sporadic human breast and ovarian cancers, the *GT198* gene is mutated in the tumor microenvironment. In breast cancer, the mutant cells are pericyte stem cells and the descendent vascular smooth muscle cell lineage, including myoepithelial cells, fibroblasts, and adipocytes (12). GT198 affects stromal cells in common solid tumors (22), including hormone-producing luteinized theca cells in ovarian cancer (23), myofibroblasts in prostate and bladder cancers (22), and pericytes in skin and brain tumors (24).

Importantly, GT198 protein is an anticancer drug target. Clinically effective oncology drugs such as paclitaxel, doxorubicin analogs, and etoposide, are found to directly bind and inhibit GT198 protein (22,25). The finding is not surprising since GT198 shares protein sequence homology with both DNA topoisomerase I and II (22), which are previously thought as targets of doxorubicin and etoposide. GT198 is also inhibited by certain herbal medicines known to be clinically successful. Since GT198 can be a drug target, targeting GT198 in immunotherapy needs investigation.

In this report, we found that GT198 is overexpressed in tumor stromal cells including pericytes, adipocytes, and myoepithelial cells in MMTV-PyMT and MMTV-Ras mouse mammary glands. We carried out vaccination using recombinant GT198 protein as antigen in MMTV-PyMT mice and confirmed that GT198 vaccine delayed mouse tumor growth. Since this is the first time to test GT198 vaccination in mice, we performed a pilot study but thoroughly evaluated each aspect in tumor development, including serum antibody production, tumor growth, lung metastasis, FACS analysis of immune cells in mouse organs, and FVB/N mouse re-implantation using GT198 positive tumor cells derived from the MMTV-PyMT tumor.

We suggest that GT198 vaccination has significant promise to warrant further investigation for human immunotherapy targeting GT198. We conclude that even though tumor initiating stimuli are distinct in human and mouse, oncoprotein GT198 is a shared target in both species resulting in a common pathway of tumor development. The oncoprotein in human breast cancer finally provides an explanation of polyomavirus-induced mammary tumor initiation in mice.

## RESULTS

### GT198 overexpression in mouse mammary gland tumor stroma

We have previously identified GT198 overexpression in human reactive breast tumor stroma due to *GT198* somatic mutations (12). Here we performed immunohistochemical staining of GT198 in mouse mammary glands of MMTV-PyMT and MMTV-Ras transgenic mice (7,8). We found robust and widespread expression of GT198 in mammary glands of transgenic but not virgin mice (**Figure 1**). The positive cells include pericytes in blood vessels, adipocytes, fibroblasts, myoepithelial cells, as well as epithelial cells. In 3-week-old MMTV-PyMT mouse mammary gland before a tumor can be detected, GT198 is already expressed in adipocytes and pericytes (**Figure 1A**). The same GT198 expression pattern is present in MMTV-Ras mice (**Figure 1B**). The expression is mostly cytoplasmic, which is a characteristic of GT198 activation with spice variant expression (21). However, tumor cells have nuclear GT198 expression which is reduced at advanced stages of the tumor (**Figure 1B**).

**Figure 1.**
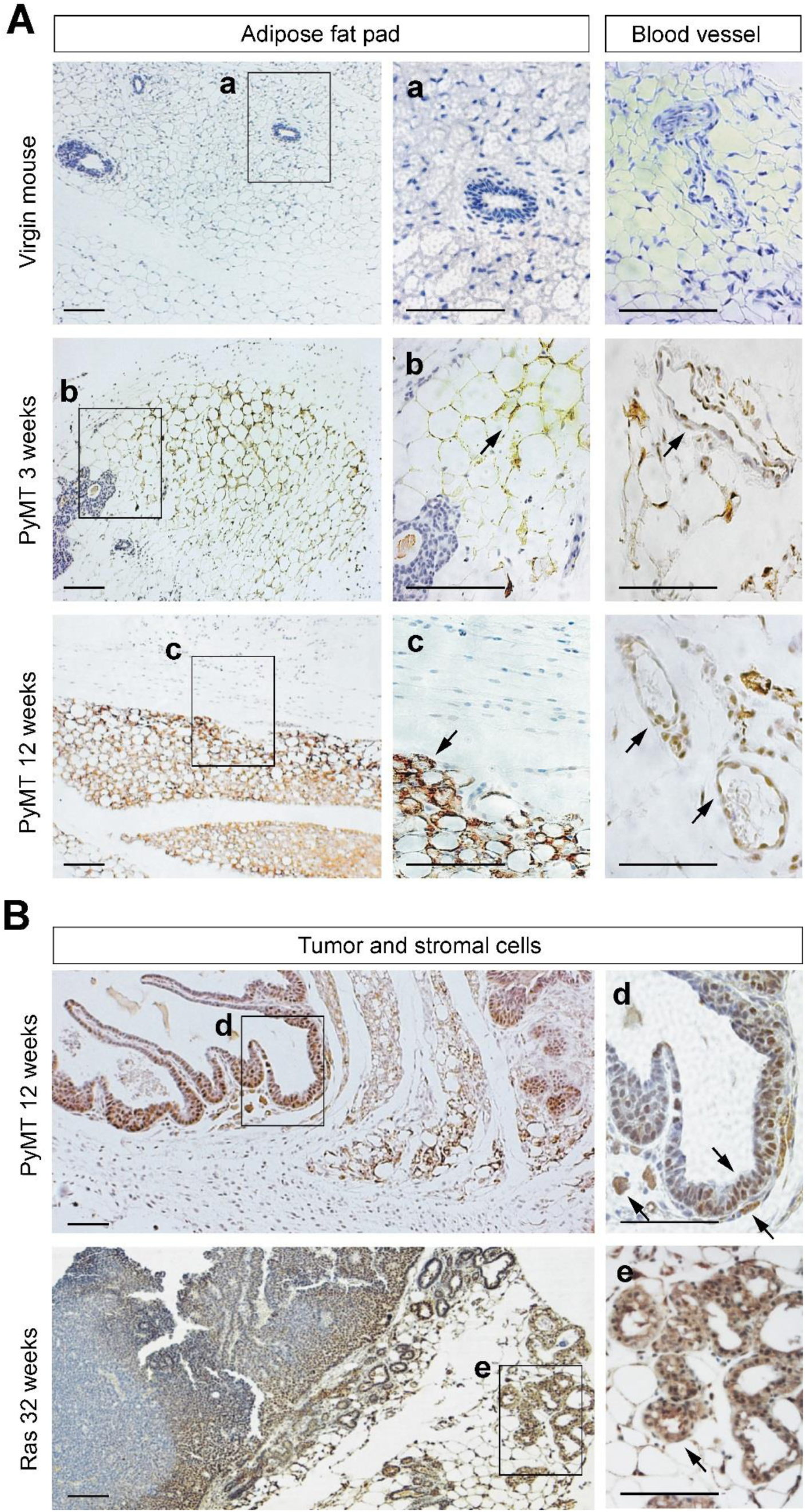
Cytoplasmic GT198 expression in mouse mammary gland tumors of MMTV-PyMT and MMTV-Ras transgenic mice. **(A)** GT198 expression in fat pads and blood vessels of mouse mammary gland tumor microenvironment. Virgin non-transgenic mouse mammary gland serves as a negative control. At three weeks of age, before tumor appears in PyMT mice, GT198 is expressed in adipocytes of the fat pad and in pericytes of the capillary. At 12 weeks when the tumor appears, cytoplasmic GT198 is highly expressed in the fat pad. **(B)** GT198 expression in mammary gland tumor of MMTV-PyMT mouse at 12 weeks of age, and in MMTV-Ras mouse at 32 weeks (8 months). Strong GT198 expression is in adipocytes, myoepithelial cells, and stromal fibroblasts. Nuclear expression is found in epithelial tumor cells at early stages of tumor. Arrows indicate GT198 positive cells. Boxed areas are enlarged at the right with corresponding labels. Sections were counter-stained with hematoxylin. Scale bars = 100 µm.

GT198 expression pattern in the mouse is similar to that in human breast cancer. However, no somatic mutation of the mouse *GT198* gene was detectable in mouse mammary glands (sequence data not shown). Because GT198 was originally cloned through activating Sos-Ras pathway (13), and GT198 variant can be stimulated by MAP kinase activity (12). The activation with overexpression of GT198 variant in transgenic mice is possibly kinase-induced and has bypassed the requirement of somatic mutations in human, which induce GT198 variant activation (21), with cytoplasmic overexpression in human breast cancers (12).

### GT198 vaccination in MMTV-PyMT mice

Oncoprotein GT198 activation is an early event in tumor development and occurs in pericytes and stromal microenvironment before tumor cells appear. We then tested GT198 vaccination to see whether serum anti-GT198 antibody in mouse has a protective effect on tumor growth.

MMTV-PyMT mice were subcutaneously vaccinated using insoluble sonicated recombinant human GST-GT198 inclusion body (n=3) or GST control (n=4) during 4-16 weeks of age in every two weeks (**Figure 2A-B** and **Supplementary Figure S1**). The adjuvant used was incomplete Freund adjuvant (IFA) due to its higher efficacy when compared to other adjuvants tested (**Supplementary Figure S1B-C).** Blood sera were collected for GT198 antibody titer analysis with tumor volumes concurrently measured at each vaccine time point. Mouse serum anti-GT198 antibody was increased in vaccinated mice than the controls, although serum antibody appeared to be consumed by GT198 positive tumors and decreased at the end of tumor development (**Figure 2B**). There was a significant delay of tumor growths in the vaccine group than control mice (**Figure 2C-D**). Importantly, tumor volumes showed a reciprocal linear correlation to serum antibody titers in all mice (**Figure 2E**). The serum anti-GT198 antibody in each mouse was further evaluated by FACS analysis, and the results suggested that vaccine-induced antibodies can recognize cell surface GT198 protein in viable cultured hypoxic AT-3 tumor cells (**Figure 2F-G**), and the increasing recognition by GT198 antibody is significant in vaccinated mice than the controls (**Figure 2H**). The antibody binding was also detectable using viable MMTV-PyMT tumor cells or PY8119 (PyMT) tumor cell line (not shown), consistent with our previous report that surface GT198 is expressed on hypoxic tumor cells (24). These data together confirmed that serum GT198 antibody would target tumor cells *in vivo* in MMTV-PyMT mice.

**Figure 2.**
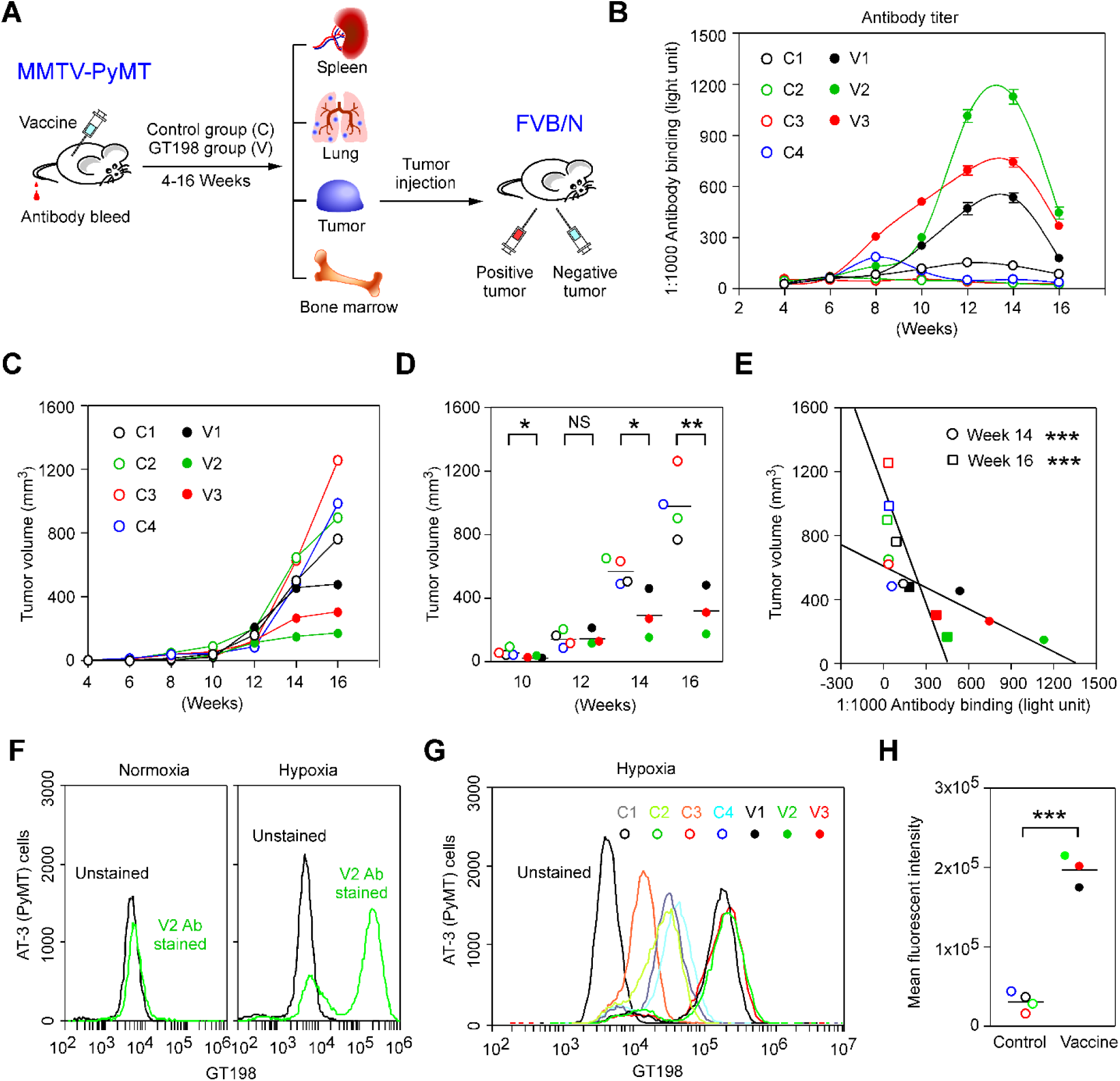
GT198 vaccination delays tumor growth in MMTV-PyMT transgenic mice. **(A)** A model of the study design. Human recombinant GST-GT198 inclusion body as antigen (100µg) was subcutaneously injected in MMTV-PyMT mice, with tumor volume measured, and tail blood collected during 4-16 weeks of age. Lung metastasis and mouse organs were analyzed at week 16. The GT198 positive and negative tumor cells were further re-implanted in pairs into each side of FVB/N mice (n=6) for tumor growth analysis. **(B)** Using his-tagged GT198 coated 96-well plates, GT198 antibody titer in diluted mouse tail blood sera (1:1000, n=2) was measured in each mouse. **(C-D)** Tumor volumes were measured and analyzed in the vaccine group (n=3) and control group (n=4). *P* * < 0.05; *P* ** < 0.01; NS, not significant. **(E)** Reciprocal linear correlation of tumor volumes with serum anti-GT198 concentrations at weeks 14 and 16. **(F)** Vaccine-induced anti-GT198 antibody (V2, week 14) increased the binding to cultured viable AT-3 (PyMT) tumor cells with PyMT origin under hypoxia condition. **(G)** GT198 antibody binding to hypoxic AT-3 cells was compared among sera from vaccine and control mice. **(H)** Same data are presented by scattergram. The vaccine group has a significant increase of anti-GT198 recognition to AT-3 tumor cells. *P* *** < 0.001.

### GT198 vaccination decreases lung metastasis

At the end of the vaccination, mouse organs including lung, spleen, bone marrow, and tumor were collected for further analyses (**Figure 2A**). The numbers of metastatic foci in both side of the lung were counted after immunohistochemical staining of GT198 (**Figure 3A**). The lung loci numbers were decreased in vaccinated mice compared to the controls (**Figure 3B-C**). Metastatic foci contain GT198 positive tumor cells, usually at the edge of foci (**Figure 3D**), which characterize the highly proliferating tumor cells (**Figure 3E**).

**Figure 3.**
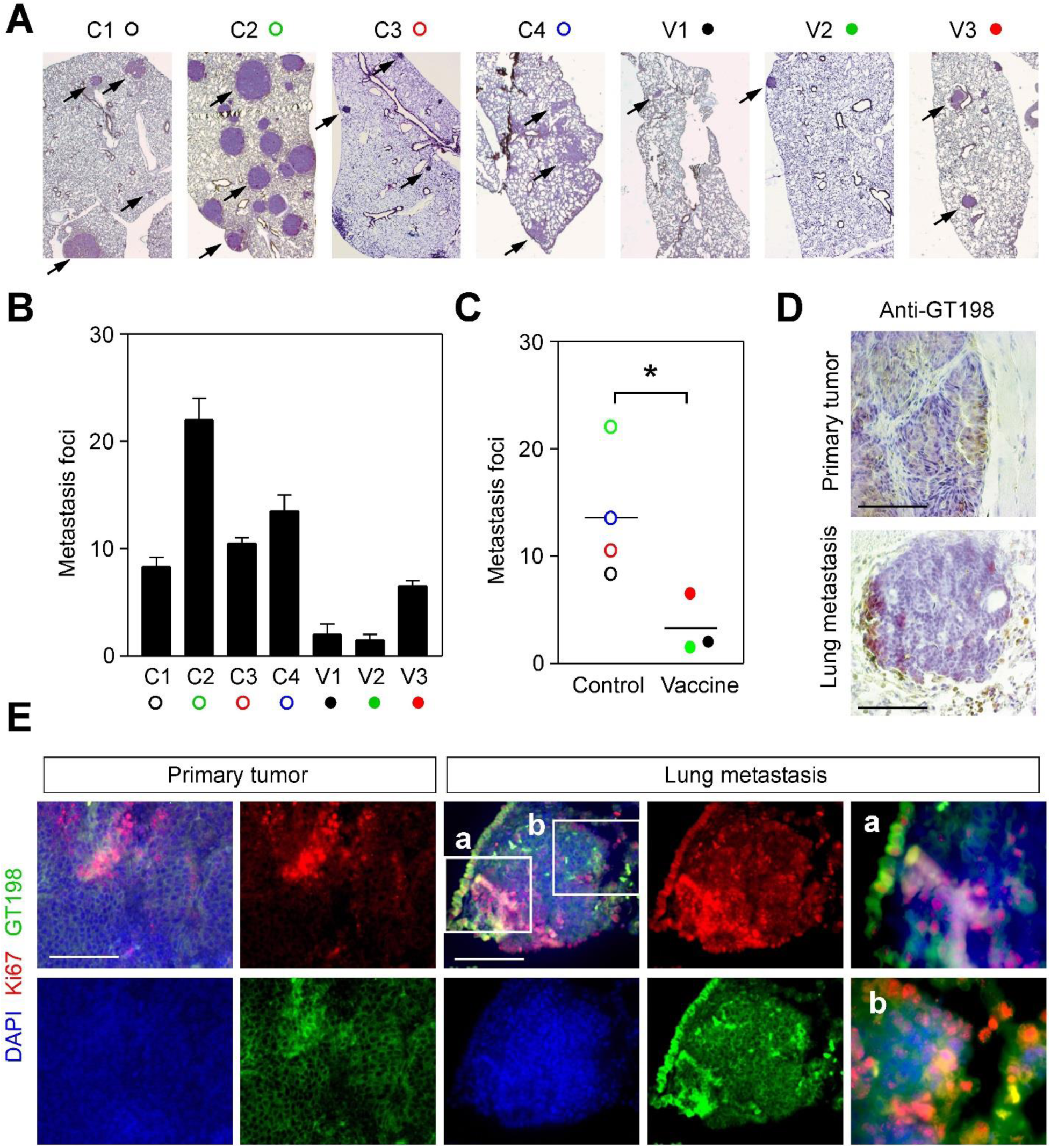
GT198 vaccination decreases tumor metastasis in the lungs of MMTV-PyMT mice. **(A)** Immunohistochemical staining of GT198 for lung metastasis in vaccine and control groups of mice. **(B)** Numbers of lung metastasis foci in each mouse. Both sides of the lung (n=2) were FFPE blocked and stained to count numbers of foci. **(C)** The averages of foci numbers are presented by scattergram. *P* * < 0.05. **(D)** Immunohistochemical staining of GT198 to compare a primary tumor in the mammary gland and a metastatic tumor in the lung. **(E)** Immunofluorescent staining of the proliferation marker Ki67 (red), showing GT198 positive cells (green) are proliferative in both primary and lung metastatic tumors. Slides were counterstained with DAPI.

### Immune cell population in vaccinated mice

We further analyzed the immune cell populations from mouse organs in representative vaccinated (V1, V2) and control mice (C1, C2). We analyzed myeloid, M1 macrophage, B cell populations (**Figure 4**). We also analyzed total lymphocyte, M2 macrophage, MDSC, T helper cells, and T cytotoxic cells (not shown). We found that there were no significant changes among vaccinated and control groups. However, there was a potential increase of myeloid, M1 macrophage, B cell populations only in tumors of vaccinated mice (**Figure 4**), although statistically insignificant. These data suggest that the mouse cellular immune response remained comparable between GT198 vaccinated mice and the control group.

**Figure 4.**
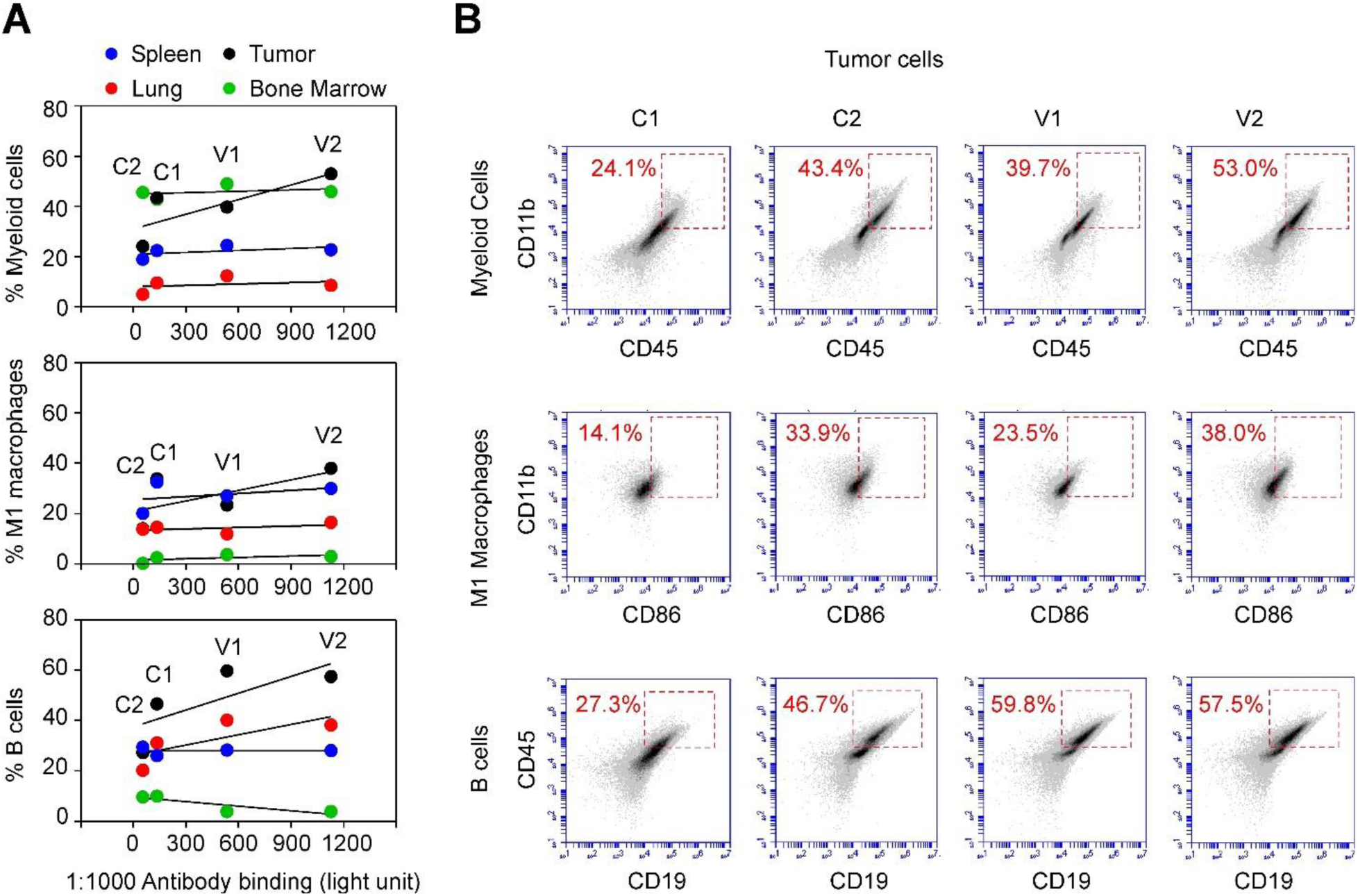
FACS analyses of immune cell populations in GT198-vaccinated and control MMTV-PyMT mice. **(A)** At week 16, FACS analyses of myeloid, M1 macrophage, and B cell populations were carried out using organs derived from the vaccine (V1, V2) and control (C1, C2) mice. Positive cell populations in the spleen (blue), lung (red), bone marrow (green), and mammary gland tumor (black) were graphed correlating to GT198 antibody titers at week 14 in each mouse. Three immune cell populations were potentially increased in tumors of mice with elevated anti-GT198 antibody, although not statistically significant. *P* > 0.05. **(B)** Flow cytometry quadrant dot plots with immune cell markers as indicated on the x-axis and y-axis. The numbers at each corner indicate the percentage of positive cells in tumors.

However, these analyses were carried out at the end of the vaccine experiment when mouse organs can be collected, and when antibody titers already dropped in the vaccine group (**Figure 2B**). It may be critical in the future to evaluate the immune response during the course of vaccination. Since there is a possibility that GT198 vaccine stimulates *in vivo* cellular immune response, in addition to detectable serum response.

### GT198 positive cells from MMTV-PyMT tumor promote tumor growth when re-implanted into non-transgenic FVB/N mice

GT198^+^ tumor stromal cells occurred before the tumor starts and GT198 vaccine delayed tumor growth. These observations suggest GT198^+^ cells may drive tumor development. To further test this hypothesis, we compared GT198^+^ and GT198^-^ cells by re-implanting them into non-transgenic FVB/N mice (n=6) with a background similar to the MMTV-PyMT mice (**Figure 5A**). We implanted both cells in each side of the same mouse to achieve better control.

**Figure 5.**
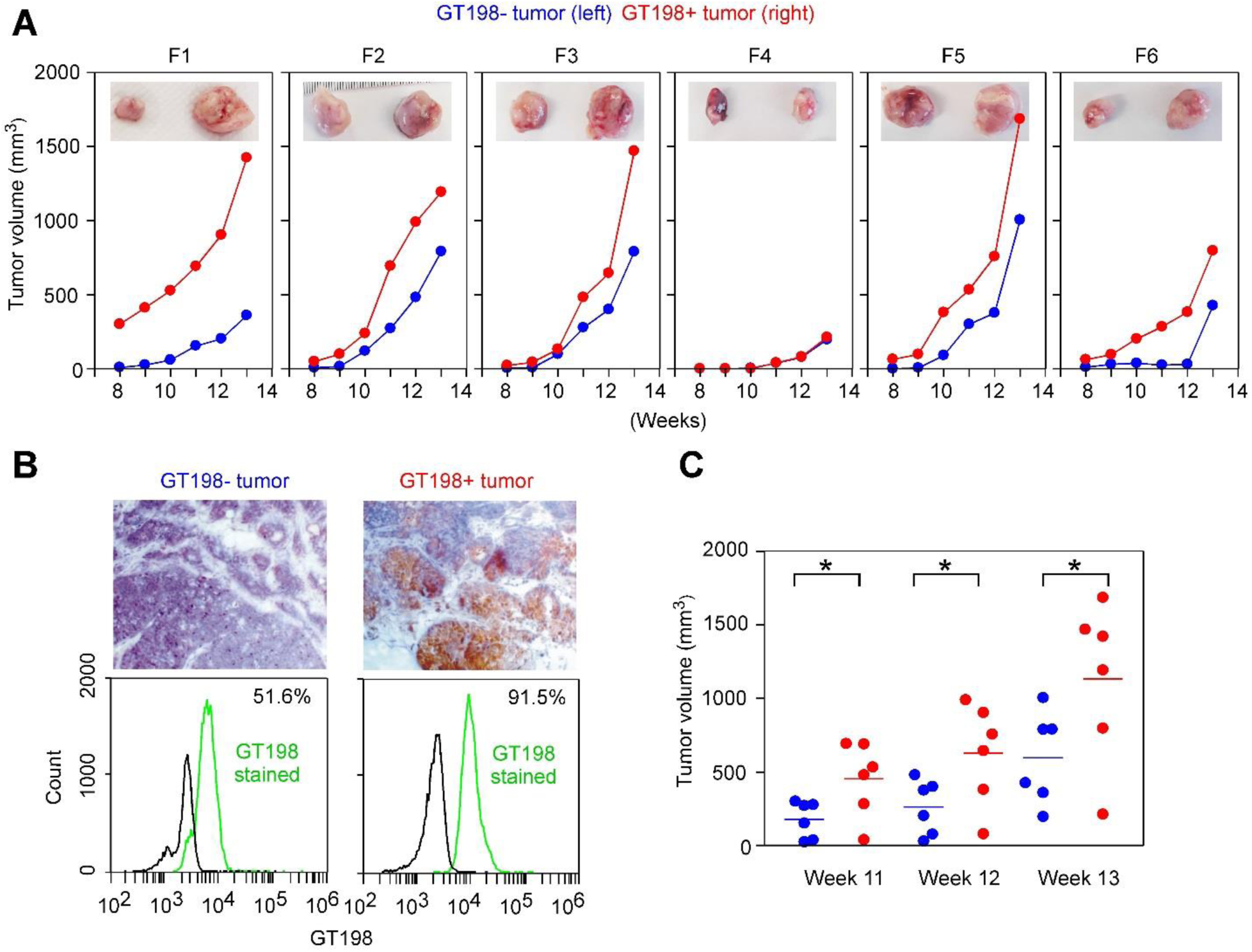
GT198-positive tumor cells promote tumor growth when re-implanted. **(A)** GT198^+^ and GT198^-^ tumor cells (10^6^ cells in 100 µl) derived from MMTV-PyMT mouse were further re-implanted into six FVB/N mice (F1-F6) with GT198^-^ cells at the left and GT198^+^ cells at the right sides of mammary glands in each mouse. The mice were followed with tumor volume measurement for additional seven weeks. The tumors at week 13 were photographed and are shown at the top (GT198^-^, left; GT198^+^, right). **(B)** Immunohistochemical staining of GT198 in MMTV-PyMT tumors used for re-implantation (top panels). FACS analysis of GT198 expression in the tumor cells. 91.5% cells from the GT198^+^ tumor are positive, although 51.6% of cells from GT198^-^ tumor are also weakly positive (bottom panels). **(C)** Tumor volumes in weeks 8-10 from (A) were analyzed by Prism scattergram showing that GT198^+^ tumor cells significantly promote tumor growth than GT198^-^ tumor cells. *P* * < 0.05.

GT198^+^ cells were derived from smaller sized MMTV-PyMT tumor, reddish in color, which contains more stromal cells confirmed by microscopy. GT198^-^ cells were derived from more massive sized tumor center, white in color, which contains mostly negative tumor cells evaluated by GT198 immunohistochemistry. These cells weakly expressed GT198 as evaluated by FACS analysis (**Figure 5B**). After an equal number of cells (10^6^ cells) were injected subcutaneously at week 6 of age in FVB/N mice, tumor volumes were measured weekly for additional 7 weeks. The tumors were collected at the end of experiments at week 13. The data showed that GT198^+^ cells significantly promoted tumor growth than GT198^-^ cells (**Figure 5A-C**). Our studies collectively suggest that GT198^+^ cells promote mouse tumor development, and GT198 vaccination decreases tumor growth in MMTV-PyMT mice.

## DISCUSSION

The polyomavirus middle T antigen-induced spontaneous mouse tumor is one of the most extensively studied mouse tumor models (4). It was predicted that the middle T antigen targets protein factors relevant to human tumorigenesis, and this is an evolutionarily conserved function utilized by the virus to propel host cell growth (2). This targeted factor has now emerged as GT198.

As a transcriptional coactivator, stem cell regulator, and crucial DNA repair factor, GT198 is in position to serve as such an oncoprotein. Deregulated GT198 activity in mutant GT198 or splice variant overexpression leads to potent apoptosis (21). We consider GT198 as a p53-like molecule in their similar dimeric structures, functions in DNA repair and cell cycle, splice variants always retaining DNA-binding domains (26), as well as unusual abundant somatic mutations (21). GT198 is also directly inhibited by paclitaxel which affects cell mitosis (25).

However, the mechanism of tumorigenesis is not merely promoting cell growth as previously thought, but is activating pericyte stem cells on blood vessels and producing stromal microenvironment responsible for epithelium growth (12). This is an important point since cytotoxic drugs are not necessarily effective antitumor drugs in human. In part because that mouse tumor cells are all fast growing and human cancer cells are slow developing in years (27).

When human cancers and rodent tumor models are compared, the GT198^+^ vessel pericytes are found as a common feature (**Figure 6**). In human breast cancer, the *GT198* gene carries somatic mutations, which activate GT198^+^ vessel pericytes in tumor microenvironment (12). In human glioblastoma xenografts of rat brain, human GT198^+^ vessel pericytes induce rat brain tumor through angiogenesis (24). In this study, the spontaneous mouse tumors carrying MMTV-PyMT or MMTV-Ras transgenes contain GT198^+^ pericytes. Similar GT198^+^ pericytes were found in MMTV-Neu transgenic mouse mammary tumors (**Supplementary Figure S2**). Consistently, distinct cancer-inducing stimuli are present in various systems, but a common GT198-mediated pathway is shared by all of them (**Figure 6**). In particular, GT198 positive angiogenic pericyte stem cells are a common feature leading to activated tumor microenvironment to propel epithelial tumor cells.

**Figure 6.**
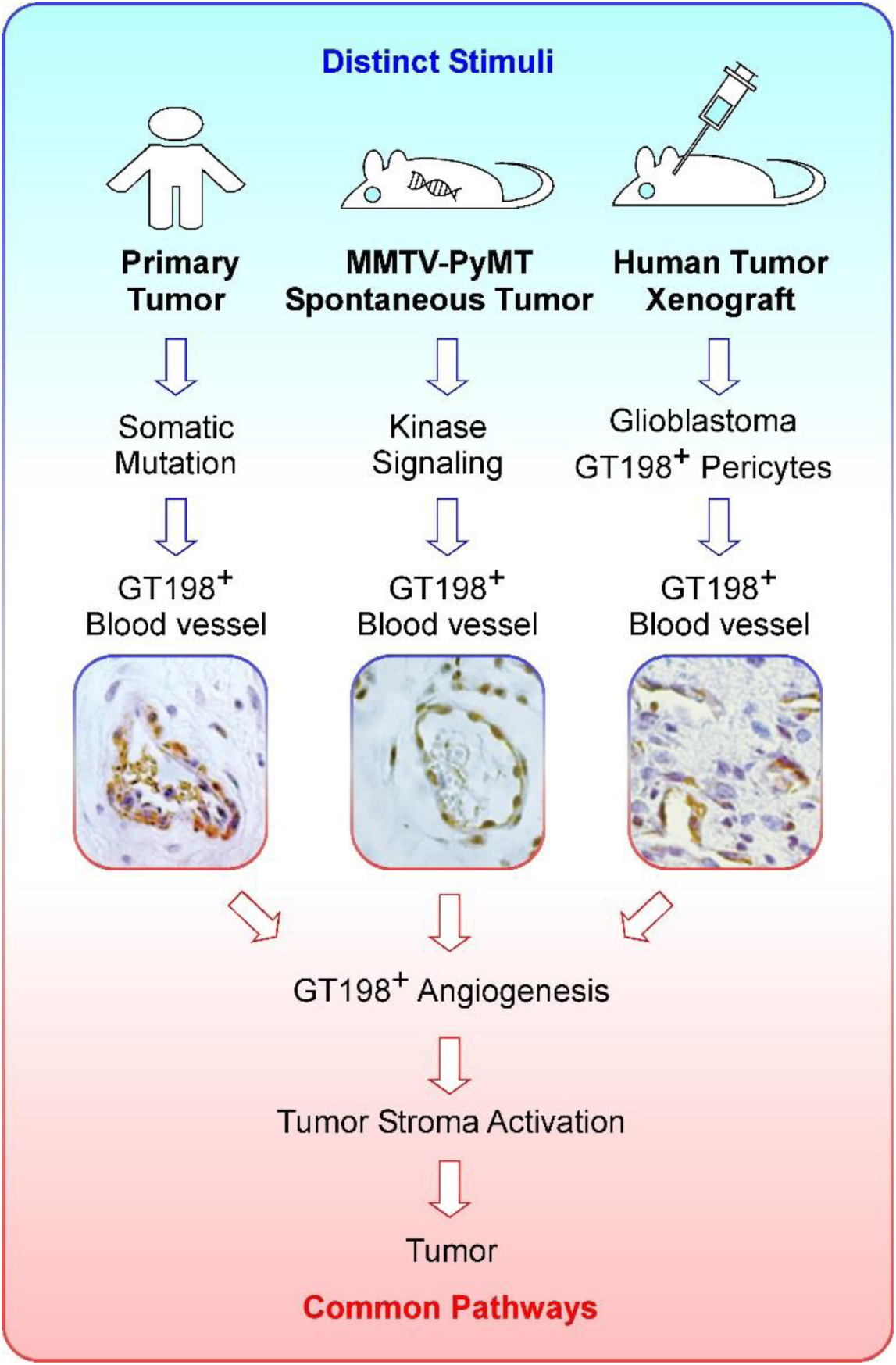
GT198^+^ pericyte stem cells are a common feature in human and mouse tumor development. GT198-mediated tumor development in human and mouse have a shared common pathway, in which pericyte stem cells in the blood vessel are activated and overexpress GT198. Activated GT198^+^ pericytes further induce tumor angiogenesis in the tumor microenvironment, and in turn, stimulate tumor growth. In human cancers (left panel), the *GT198* gene carries somatic mutations, and GT198^+^ vessel pericytes are present in multiple common solid tumors (12,22-24). A human breast cancer blood vessel is shown. In spontaneous mouse tumors carrying transgenes (middle panel), GT198^+^ pericytes are present in the blood vessel (photo from Figure 1A, PyMT 12 weeks). In human tumor xenografts (right panel), GT198^+^ vessel pericytes are activated and induce tumor angiogenesis. Human glioblastoma in rat brain is shown (24). Therefore, GT198^+^ pericyte stem cells are a shared feature in a common pathway in tumor development in both human and mouse. This model suggests that cancer initiation is distinct, but tumor development pathway is the same in both human and rodent.

The key to this mechanism is that pericytes are stem cells, and GT198 is a stem cell regulator. We have previously shown that GT198 splice variant is down-regulated during normal stem cell differentiation (**Figure 7**), so that variant activation with GT198 cytoplasmic expression in pericytes is mimicking the stem cell status or preventing them from normal differentiation. This creates a differentiation-blocked abnormal mammary gland stroma containing over activated adipocytes, fibroblasts, and myoepithelial cells; whose primary jobs are to promote the growth of epithelial cells. Similarly, GT198^+^ pericytes are present in the tumor stroma of multiple human solid tumors (24). Therefore, cancer initiation stimuli are distinct, but GT198-mediated tumor development pathway is consistent in both human and rodent (**Figure 6**).

**Figure 7.**
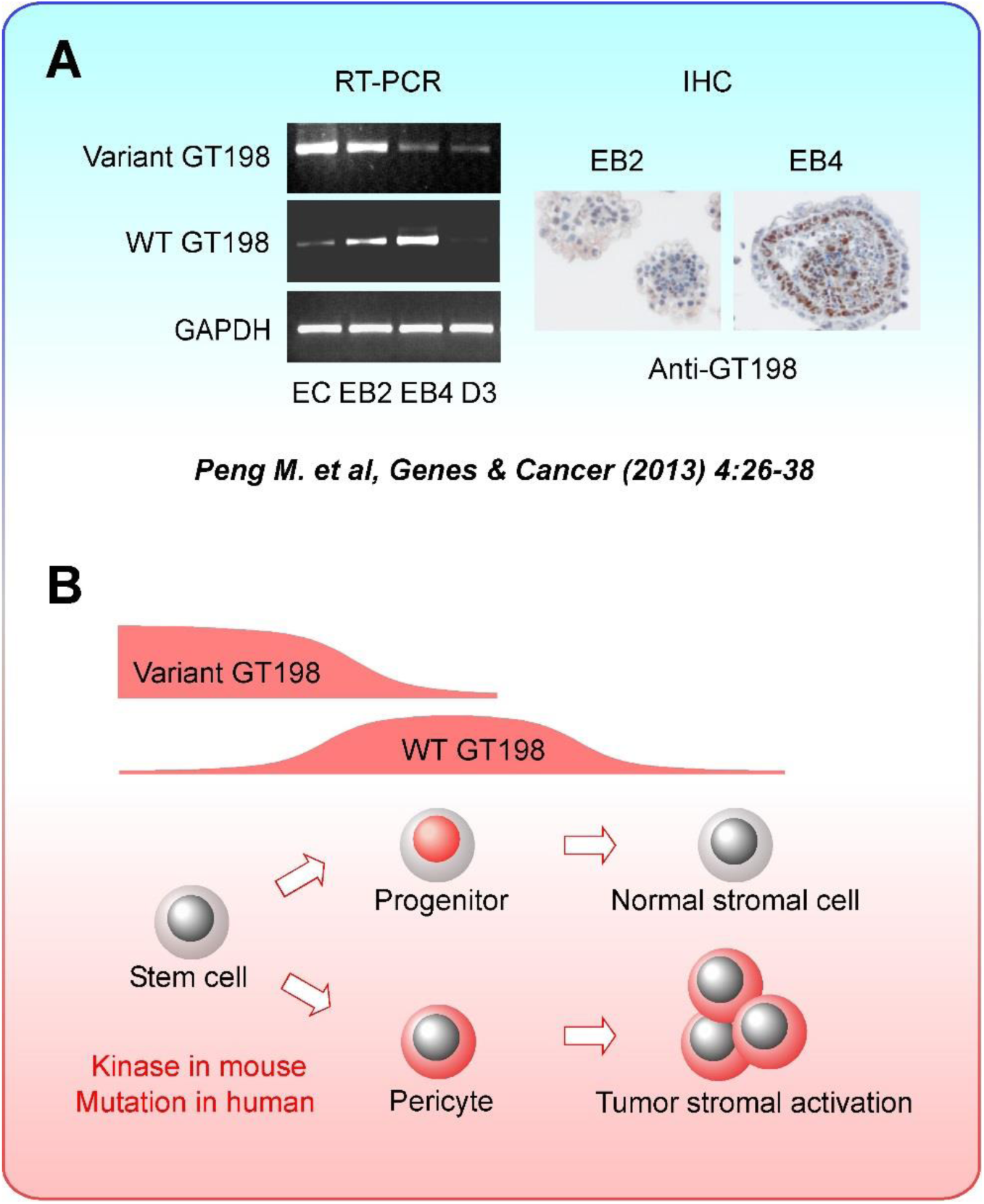
A Model of GT198 in stem cell differentiation. **(A)** GT198 regulates normal mouse embryoid body stem cell differentiation with an alternative splicing switch from its stimulatory cytoplasmic splice variant to inhibitory nuclear wild type (21). Wild type GT198 is transiently increased in mouse embryoid body, in early embryonic tissues, and is decreased in most of adult mouse tissues (24). **(B)** As a transcriptional regulator, inhibitory wild type GT198 is predicted to suppress stem cell-specific genes permitting differentiation, and is later down-regulated to release the suppression allowing differentiating genes to express. In human cancer, somatic mutations in *GT198* cause splice variant increase in breast cancer pericyte stem cells (12). In MMTV-PyMT transgenic mice, PyMT antigen induces MAP kinase pathway, which is known to stimulate variant GT198 activity (12), thereby leading to GT198 variant overexpression **(Figure 1)**. Of note, a positive feedback loop may exist between MMTV-driven PyMT and kinase-driven GT198 variant, which also stimulates the MMTV promoter (21). During mammary gland differentiation, the required downregulation of GT198 variant can be blocked by this positive feedback loop, overly activating stromal cells for tumor development.

GT198-positive cells are a driving force of tumor that can be targeted in anticancer therapy. This idea is already supported by the fact that paclitaxel and doxorubicin directly inhibit GT198 (22,25). The current study provides further support that GT198 can be a potential target in immunotherapy. However, to achieve the ideal vaccination effect, a humoral response is essential. Our data showed that all three vaccinated mice decreased their serum anti-GT198 antibody at the advanced stages of the tumor (**Figure 2B**). It appears that the serum antibodies in mice were used up due to increasing GT198^+^ tumors. Besides, we initially designed four mice in the vaccine group, but one was censored. This sick mouse had a normal amount of T cells comparable to controls but had a 50-fold decrease of B cells without production of anti-GT198, and had tumor growth outside mammary gland (not shown).

Nonetheless, it implies that GT198 vaccination may have a protective effect against tumor only when the individual is healthy with a normal hormonal response. In human, if older patients have weak immune systems, GT198 protein vaccine would be less effective, whereas monoclonal GT198 antibody therapy could be a better option. This first study of GT198 vaccination in mice aims to pave the way for the future investigations of immunotherapy targeting GT198.

In summary, we find that GT198 is overexpressed in mammary glands of MMTV-PyMT and MMTV-Ras mice. GT198 expression in mouse pericyte stem cells and tumor stromal cells are consistent with that in human reactive breast cancer stroma. GT198 vaccination in MMTV-PyMT mice delayed mouse tumor growth and reduced lung metastasis. Our study indicates that even though distinct cancer-initiation stimuli exist between mouse and human, the defective oncoprotein GT198 is a shared mechanism of tumor development in both species. Therefore, GT198 can be targeted in human cancer therapy.

## ACKNOWLEDGEMENTS

We thank Dr. Esteban Celis and Dr. Yukai He for discussion of immunotherapy. We thank Dr. Juan Wu for testing vaccine adjuvants. We thank Dr. Jianming Xu for the study of GT198 expression in mice. This work was supported in part by the Georgia Cancer Coalition Distinguished Cancer Scholar Award to L.K., the National Institutes of Health grant CA160216 and CA172048 to A.S.A; CA062130 to N.F.M.; and American Cancer Society grant IRG-14-193-01 grant to B.R.A.

---

**Supplementary Figure S1.**
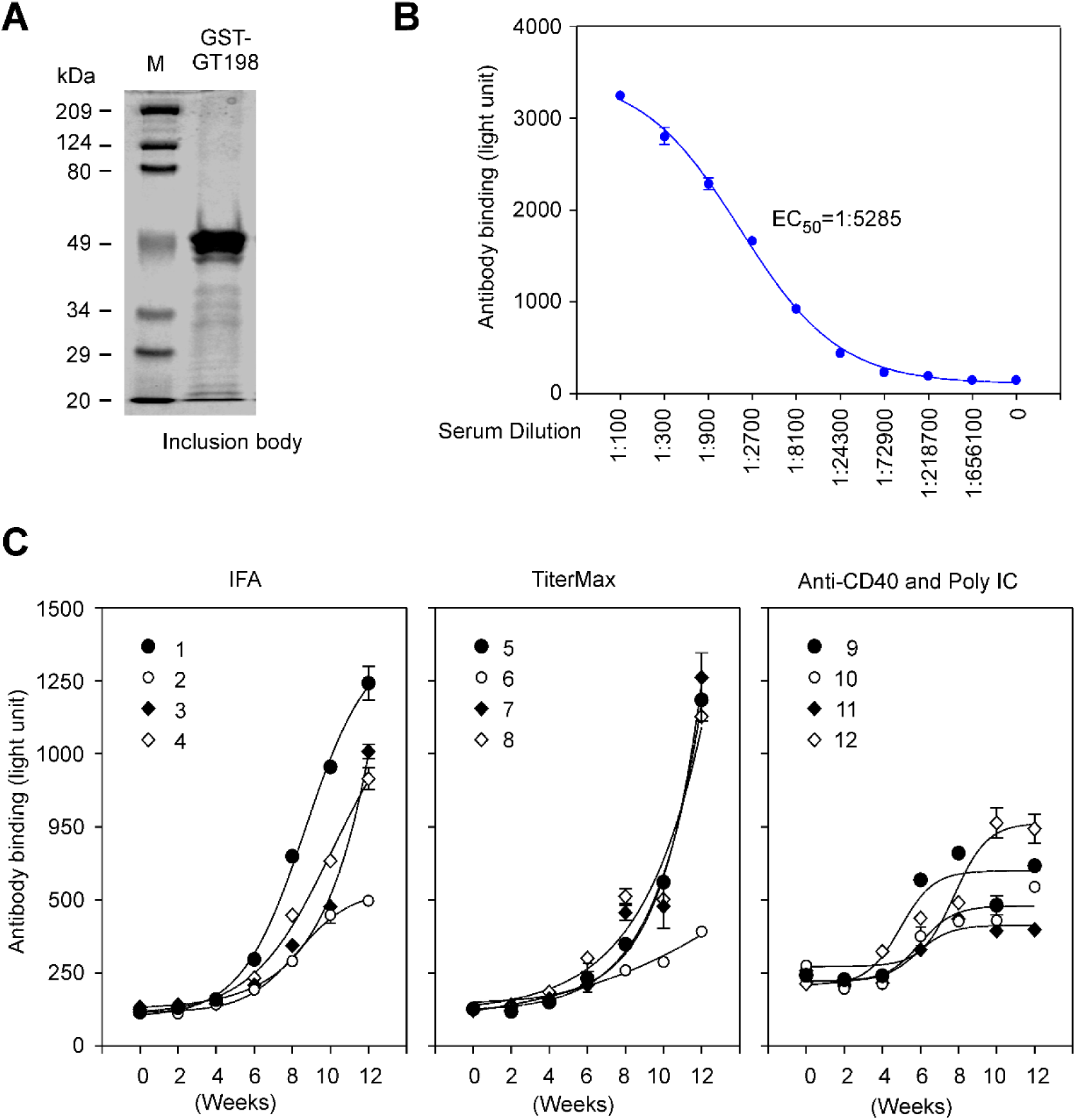
GT198 vaccine and adjuvants. (**A**) Coomassie blue staining of a protein gel showing GST-tagged GT198 fusion protein from bacteria inclusion body as antigen in vaccination. (**B**) To identify the optimized dilution in mouse sera for measuring GT198 antibody titer, GT198-vaccinated B6 mouse serum was serial diluted and tested on His-GT198 coated 96-well plates (n=2). EC50 was found as 1:5285, and thus 1:1000 and 1:3000 dilutions were chosen for the tests. Figure 2B represents the data using 1:1000 serum dilution. (**C**) To compare the adjuvants in GT198 vaccination, subcutaneous injections of Incomplete Freund’s adjuvant (IFA) or TiterMax, or intramuscular injection of Anti-CD40 and Poly IC were tested with GT198 vaccination in three groups of B6 mice (n=4). IFA yielded higher efficacy and was chosen for GT198 vaccination in MMTV-PyMT mice.

**Supplementary Figure S2.**
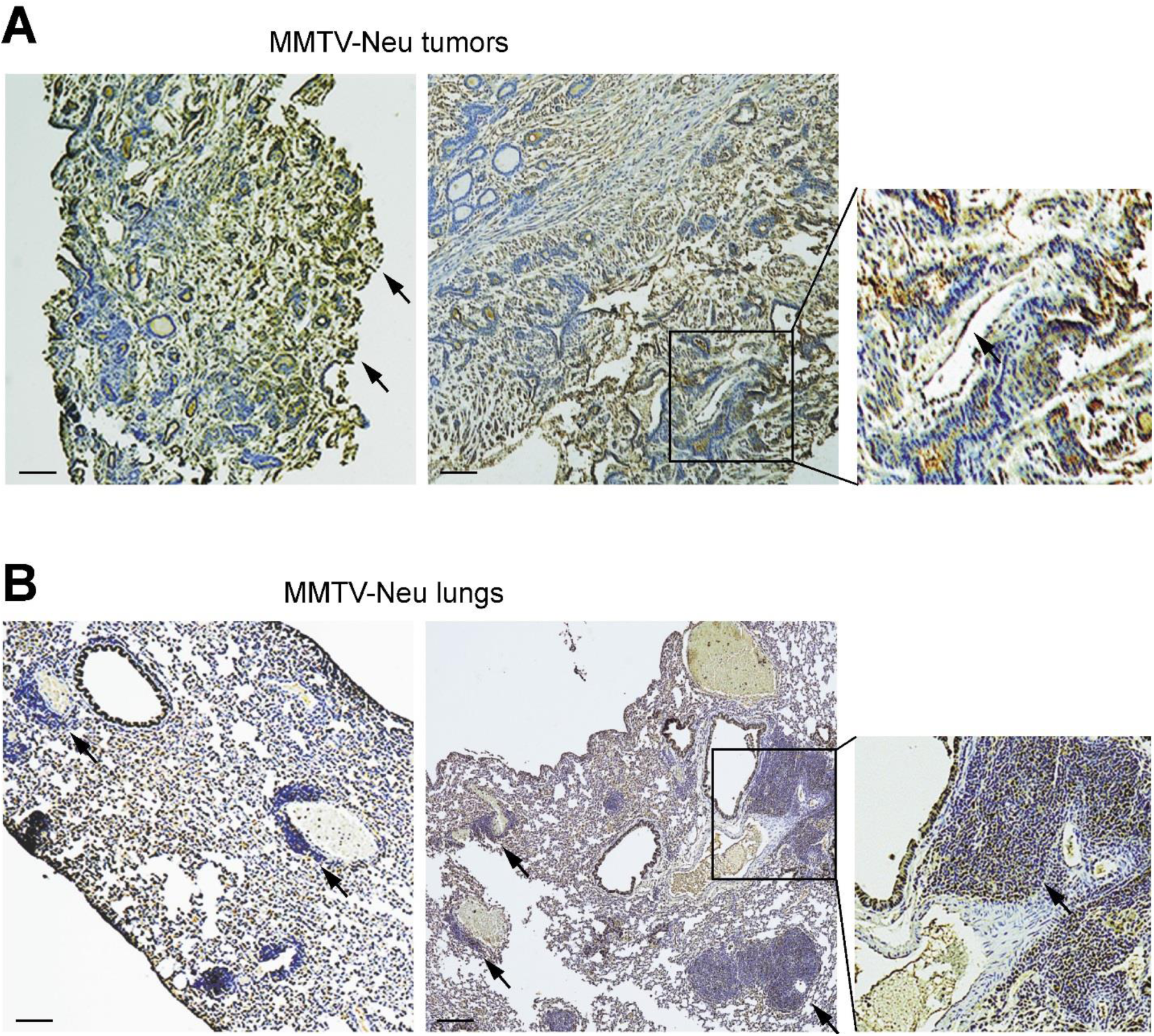
GT198 expression in MMTV-Neu mice. (**A**) Immunohistochemical staining of GT198 in mammary gland tumors of MMTV-Neu mice. Tumor stromal cells strongly express GT198 as indicated by arrows. The GT198 positive cells include blood vessel pericytes (insert). (**B**) GT198 expression in lung metastasis of MMTV-Neu mice. Lung tumor foci with GT298 expression were developed surrounding the blood vessels as indicated by arrows. Normal mouse bronchiole epithelia are known to be GT198 positive (24). Boxed areas are enlarged at the right. Sections were counter-stained with hematoxylin. Scale bars = 100 µm.

## METHODS

### Immunohistochemistry

Polyclonal rabbit antibody against GT198 was affinity purified and previously described (12,13). FFPE sections of mouse tumors and lungs from MMTV-PyMT (FVB/N-Tg(MMTV-PyVT) 634Mul/J) (7), MMTV-Ras (FVB.Cg-Tg(MMTV-vHaras)SH1Led/J) (8), and MMTV-Neu (FVB/N-Tg(MMTVneu)202Mul/J) (28), mice were deparaffinized and dehydrated through xylene and ethanol series, followed by antigen retrieval in 10 mM sodium citrate buffer, pH 6.0, containing 0.05% Triton at 90°C for 20 min. Anti-GT198 (1:200) was incubated at 4°C overnight. Antibody binding was detected using a biotinylated secondary antibody followed by detecting reagents (Abcam, Cambridge, MA). Sections were counterstained with hematoxylin.

### His-GT198 protein purification

N-terminal His-tagged recombinant human GT198 protein (aa 1-217) was expressed in *E. coli* BL21(DE3)pLysS and purified through Ni-NTA-agarose (Qiagen, #30210) as previously described (21). Proteins were eluted by 200 mM imidazole, desalted and concentrated using Amicon YM-10 spin columns before used in GT198 antibody detection.

### GT198 inclusion body as antigen in vaccination

The glutathione *S*-transferase (GST) fusion human GT198 protein was expressed in *E. coli* BL21(DE3)pLysS and insoluble inclusion body was collected as antigen in mouse vaccination since insoluble antigen has greater efficacy in vaccination. Briefly, the isolated inclusion body containing 95% pure GT198 protein (**Supplementary Figure S1A**) was repeatedly washed by sonication in PBS and sterilized by 70% ethanol. Incomplete Freund’s adjuvant (IFA) was mixed with PBS at 1:1 ratio together with GT198 protein pellet, and was sonicated using a sterilized probe to produce GT198 antigen at 1 mg/ml for each subcutaneous injection at 100 µg in 100 µl. GST was too soluble to yield inclusion body so that soluble GST protein served as control.

### GT198 vaccination in mouse tumor models

An institutional animal care and use committee (IACUC) approval was obtained (#2014-0625). In MMTV-PyMT mice, GT198 vaccinations (n=3, an additional one was censored) together with controls (n=4) were carried out in 4-week-old female mice after confirmed by genotyping. Because GT198 expression in tumor stroma can be found in as early as three weeks (**Figure 1A**), vaccination is designed to be as early as possible. Mice were subcutaneously injected at 100µg of GT198 inclusion body in 100µl IFA solution, in every other week starting from week four until week 16 (**Figure 2B**). Controls were GST in IFA. Mouse tail blood was collected at each vaccine time point to produce serum (5-10 µl). The antibody titers were measured at the end of experiment using His-tagged GT198-coated 96-well white plate (100 ng GT198 and 5 µg BSA/well), which was incubated with 200 µl of 1:1000 or 1:3000 diluted mouse sera in duplicate wells, and detected by HRP-conjugated anti-mouse antibody with ECL detection reagents. Antibody titers were counted by a Dynex luminometer. Tumor volumes were also measured at each vaccine time point from week 4 to week 16. The volume calculations were obtained using the formula V = (W(2) × L)/2 with caliper measurements. Tumor volume and GT198 antibody binding correlation were further statistically analyzed.

In FVB/N mice, tumor re-implantation was carried out in six female mice at 6-week-old of age. GT198+ tumor cells were obtained from MMTV-PyMT tumor with smaller tumor size and with tumor stromal contents. GT198-tumor cells were obtained from MMTV-PyMT tumor with larger tumor size and at the center of the tumor. The tumor cells at the center of a larger tumor are generally more GT198 negative (**Figure 1B**), in contrast to GT198 positive in tumor stroma. This was later confirmed by immunohistochemistry staining of GT198 (**Figure 5B**). Each mouse (n=6) was implanted with GT198-cells at the left, and GT198+ cells at the right sides of the mammary glands (10^6^ cells in 100 µl), and followed for additional seven weeks with tumor volume measurements.

In B6 mice for vaccine and adjuvant evaluations (n=4), GT198 vaccination using the inclusion body was similarly carried out by comparison of three adjuvants. Subcutaneous injections of Incomplete Freund’s adjuvant (IFA, 1:1) or TiterMax (1:1), or intramuscular injection of Anti-CD40 and Poly IC (50 µg each) were used with 100 µg GT198 in 100 µl volume.

### Lung metastasis analysis

Both sides of the lung in MMTV-PyMT mice were prepared in FFPE sections, immunohistochemically stained with anti-GT198 (1:150), and immunofluorescent stained with anti-GT198 and anti-Ki67 as a proliferation marker. Foci numbers were counted from both sides of the lung in the analysis.

### FACS analysis

FACS analysis and data acquisition were performed on a flow cytometer (Accuri C6, BD Biosciences). Immune cells were isolated from spleen, lung, tumor, and bone marrow from the vaccine or control MMTV-PyMT mice. FACS analysis was carried out for myeloid (CD45+, CD11b+), M1 macrophage (CD45+, CD86+, CD11b+), and B cells (CD45+, CD19+) (**Figure 4)**. Analysis for lymphocyte (CD45+), M2 macrophage (CD45+, CD206+, CD11b+), MDSC (CD45+, Gr1+, CD11b+), T helper cells (CD45+, CD4+), and T cytotoxic cells (CD45, CD8+) were also carried out but are not shown. Hypoxia AT-3 cells which is a PyMT mouse tumor cell line were generated in DMEM with 10% fetal bovine serum, in 1% oxygen and 5% CO2 at 37°C overnight. Living normoxia and hypoxia AT-3 cells were stained in PBS with 1% BSA on ice using anti-GT198 and anti-rabbit Alexa 448. A minimum of 10,000 cells within the gated region were analyzed.

### GT198 antibody titer analysis

The antibody titers were measured using collected mouse sera. Briefly, mouse tail blood (20 µl) was left on ice for 30 min and centrifuged to collect sera (10 µl). White MicroLite^TM^ 2+ 96-well plates (Thermo Scientific, #7572) were coated to dry overnight at 37°C with 400 ng/well of recombinant His-tagged GT198 proteins together with 5 µg/well of purified BSA (NEB) in a volume of 50 µl. BSA alone was included as a control for background. Duplicate wells were used for each experimental point (n = 2). The GT198-coated plates were blocked with 5% BSA in PBS with 0.1% Triton X-100 (TPBS) for 1 h. The antibody binding was carried out in 200 µl of 1:1000 or 1:3000 diluted mouse sera in duplicate wells, and detected by HRP-conjugated anti-mouse antibody with ECL detection reagents. Antibody titers were counted by a Dynex luminometer.

### Immunofluorescence

Paraffin-embedded mouse tumor tissue sections were deparaffinized through xylene and ethanol series, followed by antigen retrieval in 10 mM sodium citrate buffer, pH 6.0, containing 0.05% Triton at 90°C for 20 min. Immunofluorescence double staining was carried out in 1% horse serum using rabbit anti-GT198 (1:150) and mouse anti-Ki67 (1:200, DAKO). Secondary antibodies were anti-mouse or anti-rabbit Alexa Fluor-conjugated antibodies (Invitrogen, Carlsbad, CA). Slides were counterstained with DAPI before visualization by fluorescence microscopy.

### Statistical analysis

Statistical analyses were carried out using GraphPad Prism software. Scattergrams with means are presented using tumor volume scores, metastasis foci numbers, or fluorescent intensity. In tumor volume and antibody binding correlation assay, *P* values were determined by linear regression. In antibody titer measurements, data represent mean ± s.e.m of duplicate experiments (n = 2). Duplicates rather than triplicates were used in each experimental data point due to high consistency. *P* values in scattergrams were calculated using unpaired two-tailed t-test. * *P*<0.05, ** *P*<0.01, *** *P* <0.001; NS, not significant. A *P* value of less than 0.05 is considered statistically significant.

